# Eukaryotic initiation factor 3d Regulates Context-Dependent Pain Hypersensitivity Through the Integrated Stress Response

**DOI:** 10.64898/2025.12.21.695844

**Authors:** Subhaan M. Mian, Sera I. Nakisli, Brodie J. Woodall, Pooja J. Patel, Aariz Nackvi, Noura Elhamadany, Kree Goss, Nwasinachi Adriana Ezeji, Muhammad Saad Yousuf

## Abstract

Eukaryotic translation initiation factor 3 subunit D (eIF3d) is a noncanonical cap binding protein implicated in selective mRNA translation under stress conditions. Here, we investigate the contribution of eIF3d to pain processing using a heterozygous eIF3d knockout (eIF3d^+/−^) mouse model. We first validated this model, confirming substantial reductions in eIF3d mRNA and protein levels in dorsal root ganglia. Baseline assessments revealed no differences in mechanical, thermal, cold, or spontaneous pain behaviors between eIF3d^+/−^ (HET) and eIF3d^+/+^ (WT) mice, indicating intact basal nociceptive function. In pain models involving peripheral inflammation and metabolic stress, including methylglyoxal injection, IL-6 administration and paw incision, HET mice displayed significantly reduced mechanical and cold hypersensitivity. In contrast, HET mice exhibited increased second phase nocifensive behavior in the formalin test, possibly indicating enhanced central sensitization. Hyperalgesic priming was comparable between HET and WT mice following IL-6 exposure. Experimental autoimmune encephalomyelitis (EAE) induced mice were unaffected by eIF3d reduction. These findings demonstrate that eIF3d selectively modulates nociceptive plasticity under defined stress conditions and suggests a context dependent role in the regulation of inflammatory and central pain sensitization.

**Highlights:**

- Baseline mechanical, thermal, cold and spontaneous pain are intact in eIF3d^+/–^mice
- Methylglyoxal-evoked ISR activation and mechanical pain is blunted in eIF3d^+/–^mice
- IL-6-evoked mechanical and cold pain are reduced without altered priming
- Mechanical hypersensitivity is reduced in eIF3d^+/–^ mice with paw incision
- EAE pain is unaltered but increased pain in phase II formalin pain in eIF3d^+/–^mice

**Graphical Abstract:** 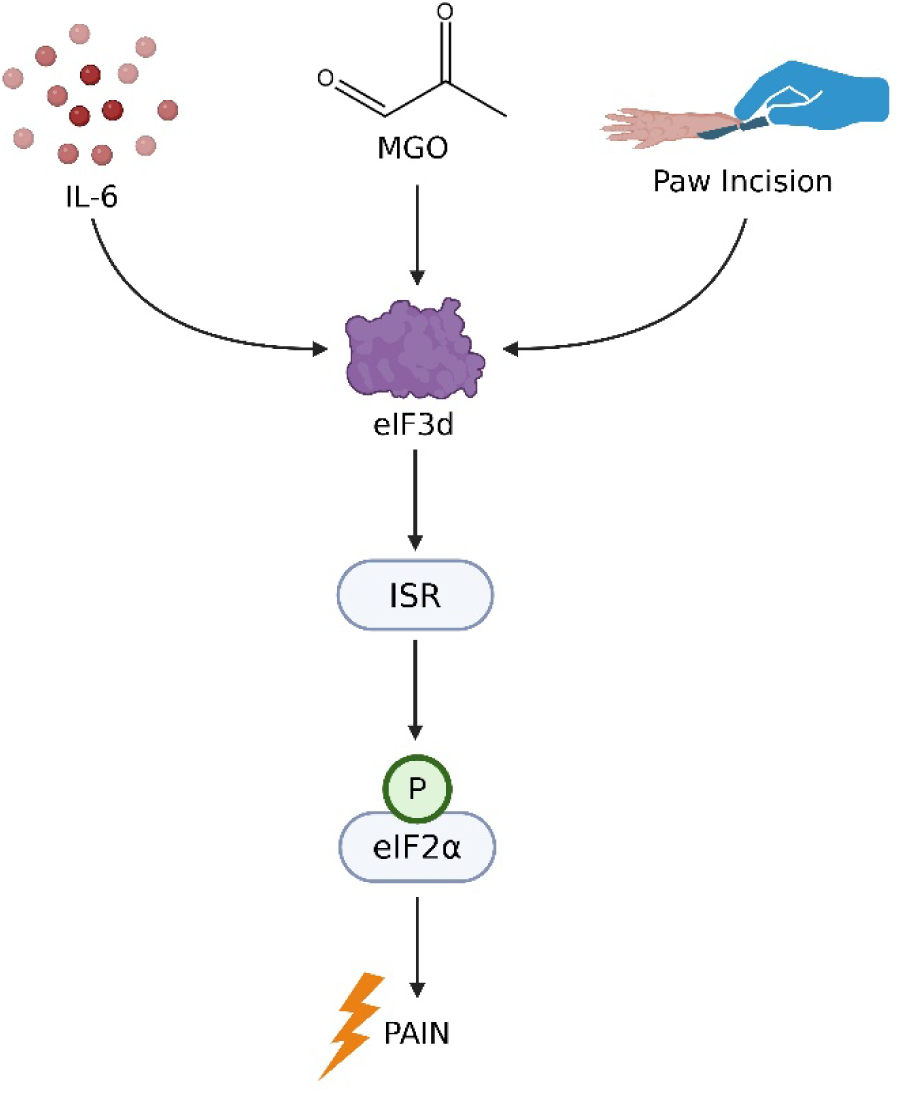

## 1. Introduction

Chronic pain remains a pervasive and debilitating condition that imposes substantial personal and societal costs [1–3]. Despite decades of research, available analgesics offer only partial relief and are frequently limited by tolerance or adverse effects, underscoring the need to define molecular mechanisms that sustain nociceptive sensitization after injury or stress. Mounting evidence indicates that translational regulation plays a pivotal role in determining how neurons and glia adapt to inflammatory, metabolic and immune challenges [4]. Rather than being governed solely by transcriptional changes, nociceptive signaling is dynamically modulated at the level of mRNA translation, enabling rapid and context-dependent adaptation to stress [4]. Central to this regulation are pathways such as the integrated stress response (ISR), which has been implicated in both neuropathic and inflammatory pain models [5–8].

Among the molecular mediators of noncanonical translation initiation is the eukaryotic initiation factor 3 (eIF3) complex. Under conditions where eIF4E-dependent translation is attenuated, eIF3 facilitates the selective translation of stress-responsive mRNAs [9].The eIF3d subunit, which possesses cap-binding activity, can enable alternative translation initiation and has been linked to ISR activation via GCN2 mediated phosphorylation of eIF2α, enhancing translation of activating transcription factor 4 (ATF4) [10–13] and inducing transcription of pro-apoptotic and pro-inflammatory genes, including JUN (c-Jun) [14, 15], the latter tightly linked to inflammation, nerve injury, and neuropathic pain [16].

Given the well-established role of altered gene expression and neuronal plasticity in chronic pain and the capacity of eIF3d to regulate translational and transcriptional networks [14, 17], we hypothesize that eIF3d influences ISR-induced pain hypersensitivity. While pharmacological inhibition of the ISR shows robust antinociceptive efficacy in animal models [5], suggesting that upstream modulation of the ISR could be therapeutically beneficial, the functional contribution of eIF3d to ISR-induced pain hypersensitivity remains insufficiently defined. Therefore, we tested whether eIF3d reduction modifies pain sensitivity across multiple pain paradigms.

Here, we describe the generation and characterization of a germline eIF3d transgenic mouse model and combined standard pain assays with sensory-neuron analyses to address how translational control gates nociception. We find that eIF3d gates context-dependent hypersensitivity: basal thresholds are intact, but mechanical and cold hypersensitivity are attenuated in metabolic (methylglyoxal), cytokine (IL-6), and surgical incision models, while neuroinflammatory EAE behaviors are unchanged and formalin Phase II is modestly increased. Mechanistically, eIF3d reduction blunts IL-6-evoked ISR activation (p-eIF2α) in mouse DRG neurons, mirroring IL-6-induced ISR in human DRG, linking eIF3d to stress-responsive translation in sensory neurons. These findings identify an eIF3d–ISR axis that selectively modulates inflammatory/metabolic pain.

## 2. Results

### 2.1 eIF3d Is Expressed in Both Human Neuronal and Non-Neuronal Cell Types, with High Expression in the DRG

Transcriptomic profiling [18] of the human *EIF3D* mRNA shows expression across various tissue types, emphasizing its relative enrichment within the human DRG (Figure 1A). Previously published spatial and single nucleus transcriptomic data [19] show that *EIF3D* is robustly expressed across all neuronal and non-neuronal subtypes within the human DRG (Figure 1B,C). Proteomic comparisons between sexes [20] indicated no significant sex-specific differences in eIF3d expression levels (Suppl. Figure 1A) and thus, subsequent analyses were not stratified by sex. Cell-type cluster annotations from the spatial and single nucleus transcriptomic dataset are detailed in Suppl. Figure 2A,B [19]. IHC analyses corroborated these transcriptomic findings. Qualitative assessment indicates consistent expression in human DRG neurons (Figure 1D, D’). Furthermore, assessments of eIF3d expression as a function of cellular diameter reveal no discernible trend suggestive of size-dependent expression variability indicated no correlation between eIF3d expression levels and cell size (Figure 1E). Correlational analysis using Pearson’s (linear) (p=0.2344) and Spearman’s (non-parametric) (p=0.3004) coefficients confirm a non-significant relationship between eIF3d expression intensity and neuronal diameter (Figure 1F). Altogether, this suggests that eIF3d expression is maintained across a broad spectrum of DRG cell morphologies.

**Figure 1.**
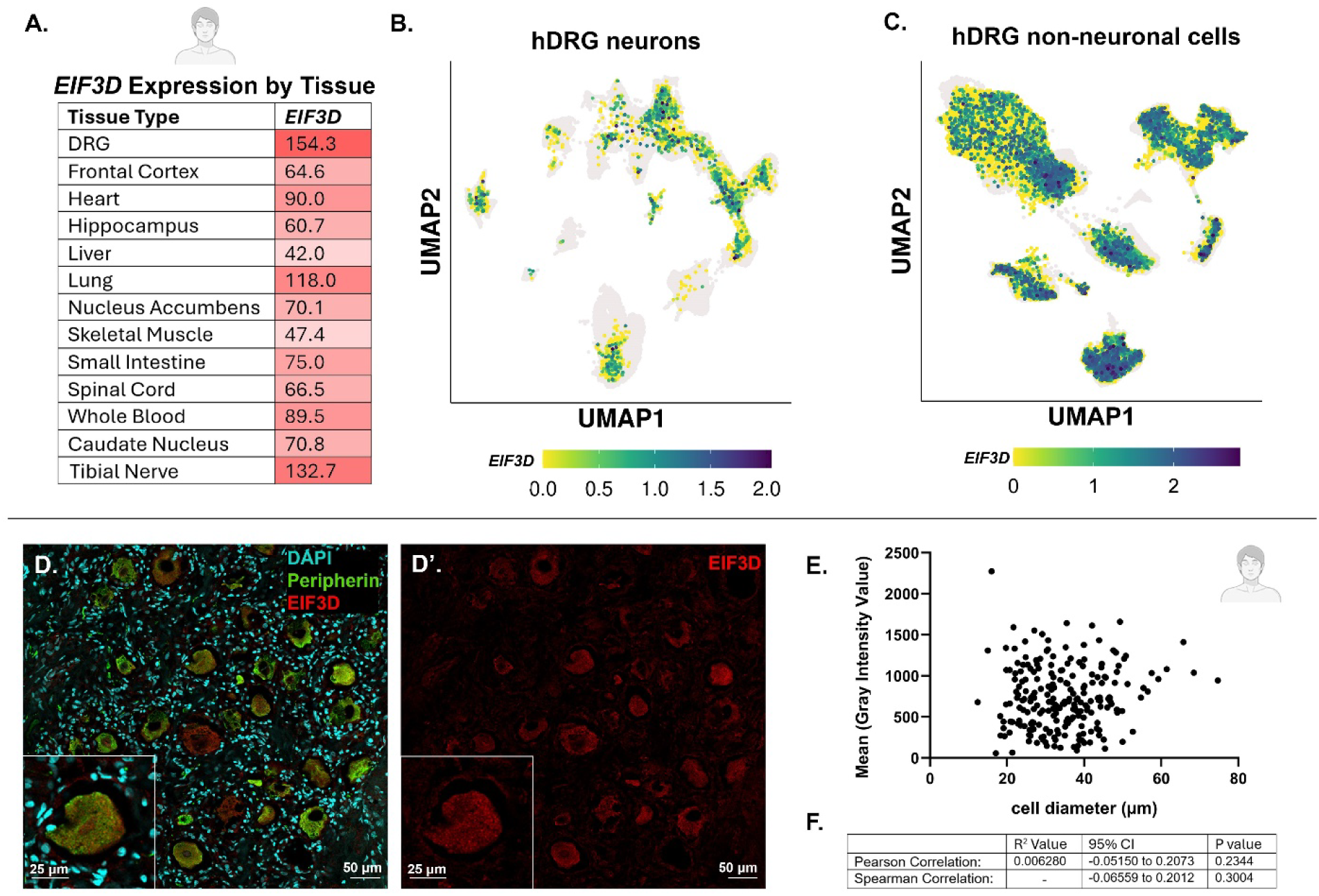
EIF3D is expressed in human DRG neurons and non-neuronal cell types, with no correlation between EIF3D expression and cell size. (A) Transcriptomic profiling shows EIF3D expression across multiple human tissue types with highest expression in the human DRG. (B, C) Spatial and single nucleus transcriptomic data illustrates EIF3D localization within human DRG neurons, neuronal subtypes (B), and non-neuronal cell types (C). (D–D′) Immunohistochemical labeling of healthy human DRG sections with DAPI (nuclei, blue), Peripherin (neuronal marker, green), and EIF3D (red) confirms neuronal expression. (E) Scatter plot depicts EIF3D protein levels do not change relative to neuronal soma size. (F) Pearson’s (linear) and Spearman’s (non-parametric) correlation analyses quantitatively show no association between EIF3D protein expression and soma size (p>0.05). *(Data from UTD CAPS Sensoryomics database)*.

### 2.2 eIF3d Expression Is Attenuated in Heterozygous Mutants of a Newly Generated Mouse Model

To investigate the in vivo function of eIF3d, we generated a germline heterozygous knockout mouse model (Figure 2A-C). Consistent with its broad representation in human DRG, eIF3d is similarly expressed across neuronal and non-neuronal populations in mouse DRG (Suppl. Figure 2 I-K).[19] Comparative spatial transcriptomic analyses across human, macaque, guinea pig and mouse demonstrate conserved eIF3d expression patterns in neuronal and non-neuronal cell types (Suppl. Fig. 2C-K)[18, 19], indicating that the translational control function of eIF3d is broadly conserved across mammals. To validate the efficacy of this model, we compared eIF3d expression levels in heterozygous (eIF3d^+/–^) knockout mice relative to wild type (eIF3d^+/+^) controls henceforth referred to as HET and WT respectively. Homozygous knockouts (eIF3d^-/-^) were not viable in utero, therefore, experiments were carried out with HET mice. RT-qPCR analysis of mouse DRG revealed an approximately 82% reduction in *EIF3d* mRNA expression in HET mice compared to WT controls (p=0.0002) (Figure 2D). ICC analysis of dissociated mouse DRG neurons and IHC analysis of mDRG tissue revealed a 47% (p<0.0001) (Figure 2E-G’) and 30% (p<0.0001) (Figure 2H-J’) reduction, respectively, in eIF3d protein in HET mice compared to WT controls. Collectively, these findings confirm that heterozygous deletion of eIF3d leads to a marked reduction in eIF3d transcript and protein abundance, thereby validating this line as a suitable in-vivo model for studying the functional role of eIF3d.

**Figure 2.**
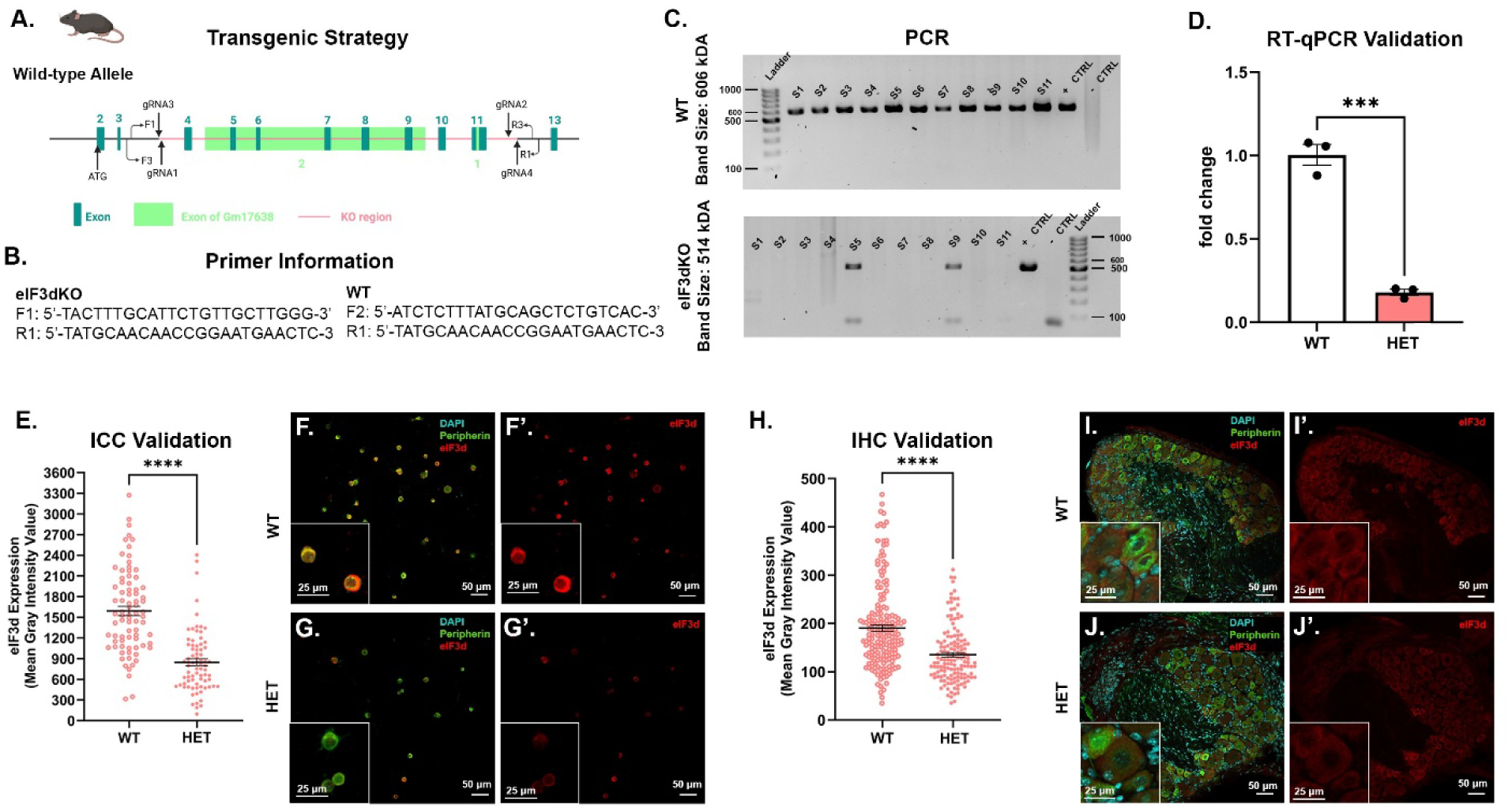
Validation of eIF3d heterozygous knockout mouse model. (A) Schematic representation of the transgenic strategy for eIF3d knockout generation, based on Cyagen’s design specifications, illustrating the targeted genomic locus, excised exons, and positions of gRNA sequences. (B) Primer sequences sizes used for genotyping PCR assays. (C) Representative PCR-based genotyping confirming eIF3d allele status. (D) Validation of heterozygous model via quantitative RT-qPCR show decrease *eIF3d* expression in HET mice (p ≤ 0.001). (E) ICC analysis of primary neuronal cultures from WT and HET mice corroborate decreased eIF3d expression in cultured cells (p<0.0001). (F, F’, G, G’) Representative images of ICC validation of the eIF3d mouse model in cultured primary mouse neurons via intensity observation. DAPI (nuclear marker, blue), Peripherin (neuronal marker, green), and eIF3d (red). (H) IHC validation of eIF3d heterozygous knockout mouse model in cryosectioned mouse DRG tissue corroborate decreased expression of eIF3d (p<0.0001). (I, I’, J, J’) Representative images for IHC validation of eIF3d expression in the mouse model via intensity observation. DAPI (nuclear marker, blue), Peripherin (neuronal marker, green), and eIF3d (red).

### 2.3 Mechanical and Thermal Nociceptive Stimuli Exhibit Comparable Pain Behaviors in eIF3d reduced Mice Compared to WT Controls

Having validated the genetic model, we next examined whether reduced eIF3d expression impacts basal nociceptive sensitivity. To determine whether partial loss of eIF3d expression impacts baseline nociceptive thresholds, we conducted behavioral assessments across multiple pain modalities in naive mice: mechanical sensitivity was tested using von Frey filaments, cold allodynia was assessed via the acetone evaporation assay, thermal nociception was measured using the Hargreaves test and spontaneous pain behavior was evaluated using the Mouse Grimace Scale. Across all modalities, HET mice exhibited no significant differences in behavioral responses compared to WT controls. Withdrawal thresholds and latencies in von Frey, acetone and Hargreaves tests were comparable between genotypes (von Frey, p=0.9629, unpaired t-test; Paw Withdrawal, p=0.8424, unpaired t-test; Acetone, p=0.9895, unpaired t-test) (Suppl. Figure 3A-C) and no evidence of spontaneous grimacing was observed in the HET mice under baseline conditions (Suppl. Figure 3D). These findings indicate that a reduction in eIF3d expression does not affect baseline pain processing, suggesting that eIF3d is not essential for normal nociceptive function in the absence of injury or inflammation.

### 2.4 Reduction of eIF3d Decreases Methylglyoxal Induced Mechanical Hypersensitivity in Mice

We next sought to determine whether eIF3d modulates nociception under metabolic stress using methylglyoxal (MGO), a dicarbonyl metabolite linked to diabetic neuropathy that triggers ISR-dependent pain hypersensitivity [21–24] and tested whether partial eIF3d loss prevents MGO induced pain. MGO has been closely associated with elevated reactive oxygen species (ROS) production and an associated disruption of cellular antioxidant defenses, contributing to oxidative stress, hyperglycemia, inflammation and aging. [22] One of MGO’s pathological effects involves the formation of advanced glycation end products (AGEs), which modify proteins and lipids, impair neuronal function and promote axonal injury, all hallmarks of neuropathic pain. [22, 24] Prior studies have demonstrated that elevated MGO levels activate the ISR and induce mechanical hypersensitivity in rodent models [5, 6].To assess the contribution of eIF3d to MGO induced pain, we administered 20 ng MGO intraplantarly, a dose previously shown to elicit ISR-mediated hypersensitivity [6]. MGO injection induced mechanical hypersensitivity in HET mice compared to WT controls (Figure 3A), as assessed by von Frey (genotype: F(1, 8)=21.99, p=0.0016; two-way ANOVA) over the 0-120 hour post-injection period (genotype x time: F (6, 48)=7.972, P<0.0001, Two-way ANOVA). Šídák’s multiple comparisons test identified significant differences between HET and WT mice at 48 hours (p<0.0001), 72 hours (p=0.0003) and 96 hours (p=0.0122) post MGO injection (Figure 3A). Significantly higher von Frey thresholds that the HETs exhibited when compared to WT controls beginning at 48 hours post injection, indicate decreased mechanical pain sensitivity in HET mice. This reduced sensitivity persisted through 72 and 96 hours, with thresholds returning to WT levels by 120 hours. These results suggest that partial loss of eIF3d reduces MGO induced mechanical hypersensitivity, implicating eIF3d as a potential modulator of neuropathic pain via regulation of stress responsive translational pathways.

**Figure 3.**
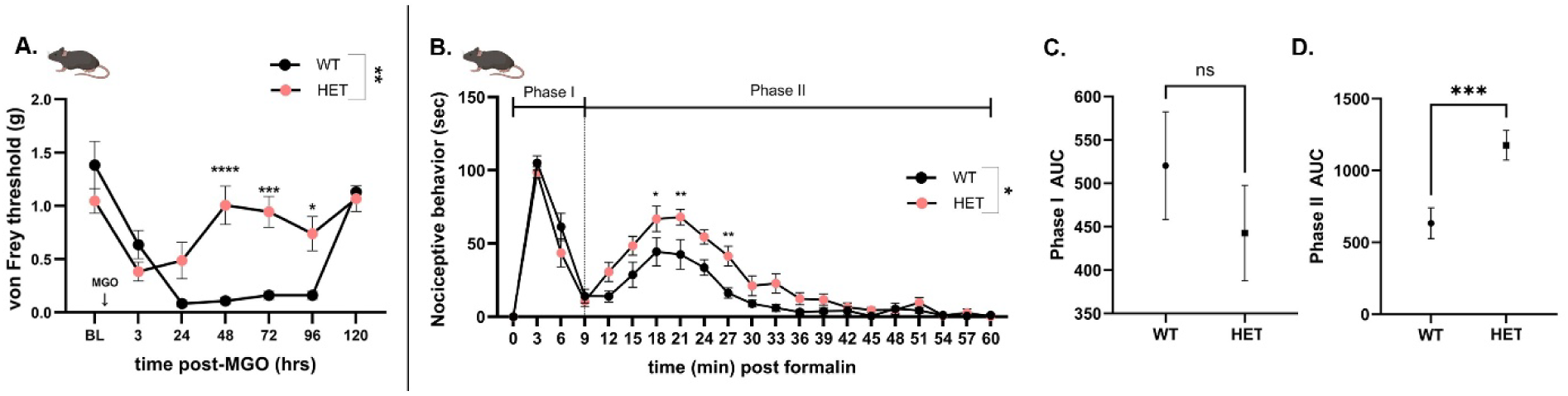
Decreased eIF3d reduces MGO induced pain and accelerates recovery from formalin evoked pain in mice. (A) MGO injection induced mechanical hypersensitivity in HET mice compared to WT controls, as assessed by von Frey (g) over the 0-120 hour post-injection period. Significant differences between HET and WT mice were identified at 48 hours (p<0.0001), 72 hours (p≤0.001), and 96 hours (p≤0.05) post MGO injection with a significant difference between genotypes (p≤0.01). (B) Overall nociceptive behavior showed a significant difference between HET and WT mice over the 0-60 minutes post formalin period (p≤0.05). There was no difference identified between WT and HET mice during Phase I of the formalin test (p>0.05). However, a significant increase in pain behavior was observed in HET mice during Phase II at 18 min (p≤0.05), 21 min (p≤0.01) and 27 min (p≤0.01). (C) Area under the curve (AUC) of Phase I for nociceptive behavior in WT and HET mice showed no significance following formalin induced pain (p>0.05). (D) AUC of Phase II for nociceptive behavior in showed significant increase in HET mice compared to WT mice following formalin induced pain (p≤0.001).

### 2.5 Loss of eIF3d in Mice Exacerbates Second Phase Formalin Evoked Nocifensive Behaviors

To investigate the role of eIF3d in inflammatory pain, we employed the formalin test, which induces a biphasic nociceptive response reflective of distinct underlying mechanisms. The initial phase (Phase I), occurring within the first 10 minutes post injection, reflects acute activation of peripheral nociceptors. This is followed by a prolonged second phase (Phase II), which is thought to arise from central sensitization mechanisms within the spinal cord, including inflammatory and synaptic plasticity related processes. To assess whether eIF3d modulates acute or persistent nociceptive responses, we performed the formalin test in both WT and HET mice. Overall nociceptive behavior showed a significant difference between HET and WT mice (Figure 3B) (genotype: F (1, 11)=7.542, p=0.0190; two-way ANOVA) over the 0-60 minutes post formalin period (genotype x time: F (20, 220)=3.015, P<0.0001, Two-way ANOVA). Šídák’s multiple comparisons test identified no difference between WT and HET mice during Phase I of the formalin test however a significant increase in pain behavior was observed in HET mice during Phase II at 18 min(p=0.0308), 21 min (p=0.0082) and 27 min (p=0.0084) (Figure 3B). However specifically during Phase I, no significant differences in nocifensive behaviors (measured by shaking, paw licking and lifting) were observed between genotypes as identified by area under the curve (AUC) analysis of Phase I (p=0.3538, Welch’s t-test) (Figure 3C), suggesting that peripheral nociceptor activation remains intact in HET mice. However, AUC analysis for Phase II show that HET mice exhibited significantly elevated nocifensive behavior compared to WT controls following formalin induced pain (p=0.0003, Welch’s t-test) (Figure 3D), indicating an enhanced central sensitization component. These findings suggest that partial loss of eIF3d may dysregulate spinal mechanisms involved in the maintenance or amplification of inflammatory pain.

### 2.6 IL-6 Leads to Attenuated Mechanical and Cold Hypersensitivity in eIF3d-Deficient Mice Compared to WT Controls While Maintaining Hyperalgesic Priming

Given the role of cytokines in modulating pain states [25, 26], we next evaluated whether eIF3d influences responses to interleukin-6 (IL-6), a key mediator of inflammatory hypersensitivity [27]. To investigate the contribution of eIF3d to cytokine induced nociceptive plasticity, we administered intraplantar injections of recombinant IL-6 at two concentrations (0.2 ng and 0.5 ng in 10 µL) to WT and HET mice. Mechanical sensitivity was assessed using von Frey and cold sensitivity was measured using the acetone test. Intraplantar (IPl) injection of 0.2 ng IL-6 significantly altered mechanical hypersensitivity in HET compared to WT mice (genotype x time: F (6, 66)=3.108, p=0.0096; two-way ANOVA). HET mice began exhibiting reduced mechanical hypersensitivity, as observed by increased von Frey thresholds (g). relative to WT mice, emerging at 6 hours post-injection (Figure 4A) (p=0.1046, Šídák’s multiple comparisons) and becoming significantly increased at 24 hours post 0.2ng injection (p=0.0455, Šídák’s multiple comparisons). An IPl injection of 0.5 ng IL-6 significantly altered mechanical hypersensitivity in HET mice when compared to WT (Figure 4E) (genotype: F (1, 12)=5.043, p=0.0443; two-way ANOVA). By 6 hours post injection, HET mice experienced a significant reduction in mechanical hypersensitivity (p=0.0467, Šídák’s multiple comparisons). Together these results indicate a decrease mechanical hypersensitivity response with reduced eIF3d in mice.

**Figure 4.**
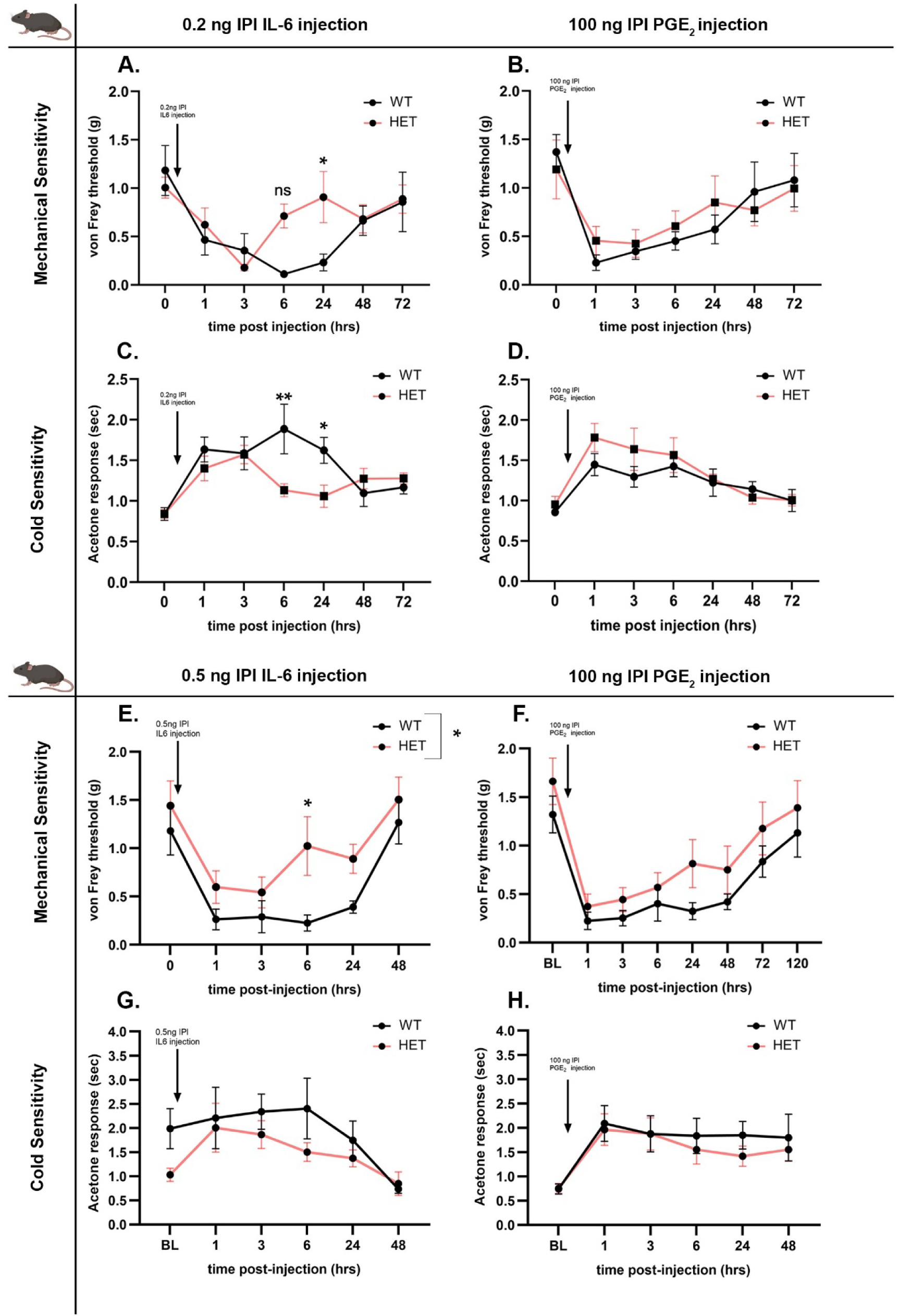
IL-6 Leads to Attenuated Mechanical and Cold Hypersensitivity in eIF3d-Deficient Mice Compared to WT Controls While Maintaining Hyperalgesic Priming. (A) Intraplantar (IPl) injection of 0.2 ng IL-6 significantly reduced mechanical hypersensitivity in HET mice compared to WT controls at 24 hours post-injection (p≤0.05). (B) To assess hyperalgesic priming for mechanical hypersensitivity, 100 ng of PGE_2_ was administered via IPl injection. Hyperalgesic priming was unchanged at any time point post-injection in either HET or WT mice (p>0.05). (C) IPl injection of 0.2 ng IL-6 significantly reduced cold hypersensitivity in HET mice compared to WT mice over a 72-hour observation period (p≤0.01). HET mice exhibit a significant reduction in cold hypersensitivity compared to WT controls at 6 hours (p≤0.01) and 24 hours (p≤0.05) post-injection. (D)100 ng of PGE_2_ was administered via IPl injection to assess hyperalgesic priming for cold hypersensitivity. Hyperalgesic priming was unchanged at any time point post-injection in either HET or WT mice (p>0.05). (E) IPl injection of 0.5 ng IL-6 significantly reduced in mechanical hypersensitivity in HET mice compared to WT controls at 6 hours post-injection (p≤0.05) with a significant difference between genotypes (p≤0.05). (F) 100 ng of PGE_2_ was administered via IPl injection to assess hyperalgesic priming for mechanical hypersensitivity. Hyperalgesic priming was unchanged at any time point post-injection in either HET or WT mice (p>0.05). (G) IPl injection of 0.5 ng IL-6 did not significantly alter cold hypersensitivity in either HET or WT mice over a 48-hour observation period (p>0.05). However, the effect profile qualitatively mirrored the trend seen with 0.2 ng IL-6. (H)100 ng of PGE_2_ was administered via IPl injection to assess hyperalgesic priming for cold hypersensitivity. Hyperalgesic priming was unchanged at any time point post-injection in either HET or WT mice (p>0.05).

Similarly, IPl injection of 0.2 ng IL-6 significantly reduced cold hypersensitivity in HET mice compared to WT mice (Figure 4C) over a 72-hour observation period (genotype x time: F (6, 66)=4.151, p=0.0013; two-way ANOVA). HET mice displayed significantly reduced nocifensive response by 6 hours (p=0.0026, Šídák’s multiple comparisons) and continuing to 24 hours (p=0.0462, Šídák’s multiple comparisons) in the acetone assay following the 0.2 ng IL-6 injection, with a similar but nonsignificant trend observed at the 0.5 ng dose (Figure 4G) (genotype x time: F (5, 60)=1.028, p=0.4092; two-way ANOVA. Both the mechanical and cold hypersensitivity in all groups had resolved by 48 hours post injection.

To determine whether eIF3d influences the persistence of sensitization, we assessed hyperalgesic priming. An injection of prostaglandin E_2_ (PGE_2_; 100 ng), at a subthreshold concentration known not to evoke hypersensitivity in naive animals, was administered three days following resolution of IL-6-induced hypersensitivity. WT and HET mice showed no change in mechanical sensitivity in response to 100 ng PGE_2_ after 0.2ng (Figure 4B) (genotype: F(1,77)=0.1374, p=0.7119; genotype x time: F(6,77)=0.4470, p=0.8450; two-way ANOVA) and 0.5ng (Figure 4F) (genotype: F (1, 12)=2.171, p=0.1664; genotype x time: F (7, 84)=0.2562, p=0.9688; two-way ANOVA) IL-6 treatment. Likewise, an injection of 100 ng PGE_2_ evoked comparable cold sensitivity in both WT and HET mice at both IL6 injections of 0.2ng (Figure 4D) (genotype: :F (1, 77), p=0.1183; genotype x time: F (6, 77)=0.6590, p=0.6828; two-way ANOVA) and 0.5 ng (Figure 4H) (genotype: F (1, 77)=0.9357, p=0.3364; genotype x time: F (6, 77)=0.1995, p=0.9760; two-way ANOVA). Suggesting that hyperalgesic priming was not affected in HET mice when compared to WT controls across high and low dose of IL-6 administration. These findings collectively suggest that eIF3d contributes to acute cytokine mediated hypersensitivity but priming is not affected by reduce eIF3d levels.

### 2.7 eIF3d modulates IL-6 induced Integrated Stress response activation in DRG neurons

p-eIF2α is a well established marker of integrated stress response (ISR) activation. Upon activation, ISR leads to increased p-eIF2α levels, which in turn suppresses global protein synthesis while promoting the translation of stress-adaptive proteins. [8, 28] This translational reprogramming can contribute to neuronal plasticity and pain development. Recent studies place eIF3d upstream of the ISR: under metabolic stress, hypophosphorylated eIF3d (near its cap-binding pocket) engages noncanonical cap binding to redirect translation toward stress-response and glucose-homeostasis mRNAs [29]; during persistent stress, eIF3d promotes GCN2 translation to increase eIF2α phosphorylation and upregulates the m6A demethylase ALKBH5 to drive 5′UTR demethylation of ISR targets (including ATF4), thereby amplifying ATF4 translation and ISR output [11]. Given that IL-6 is known to induce pain related behaviors, we sought to determine whether IL-6 drives ISR activation in the DRG neurons. To assess this, we cultured primary DRG neurons from WT and HET mice and treated them with either IL-6 or vehicle. ICC for p-eIF2α revealed that IL-6 treatment significantly increased p-eIF2α levels in WT DRG neurons compared to vehicle treated WT controls (Figure 5A,B-C’) (p=0.0020, two-way ANOVA), indicating robust ISR activation in response to IL-6. In contrast, HET DRG neurons did not exhibit a significant increase in p-eIF2α following IL-6 treatment relative to their vehicle treated counterparts (Figure 5A,D-E’) (p=0.1940, two-way ANOVA). p-eIF2α expression in IL-6 treated HET neurons was markedly lower than both vehicle (Figure 5A,B,B’) (p=0.0003, two-way ANOVA) and IL-6 treated WT neurons (Figure 5A,C,C’) (p=<0.0001, two-way ANOVA). To extend these findings to humans, we cultured healthy human DRG neurons and treated them with IL-6 or vehicle. IHC analysis demonstrated a significant increase in p-eIF2α expression following IL-6 exposure (Figure 5F,G-H’) (p=0.0008, unpaired t-test), mirroring the response observed in WT mouse neurons and suggesting an ISR activation. These results suggest that IL-6 activates the ISR in both mouse and human sensory neurons and that eIF3d is necessary for this IL-6 driven p-eIF2α response. Reduced eIF3d expression blunts ISR activation, has a potential role in modulating inflammation induced pain signaling.

**Figure 5.**
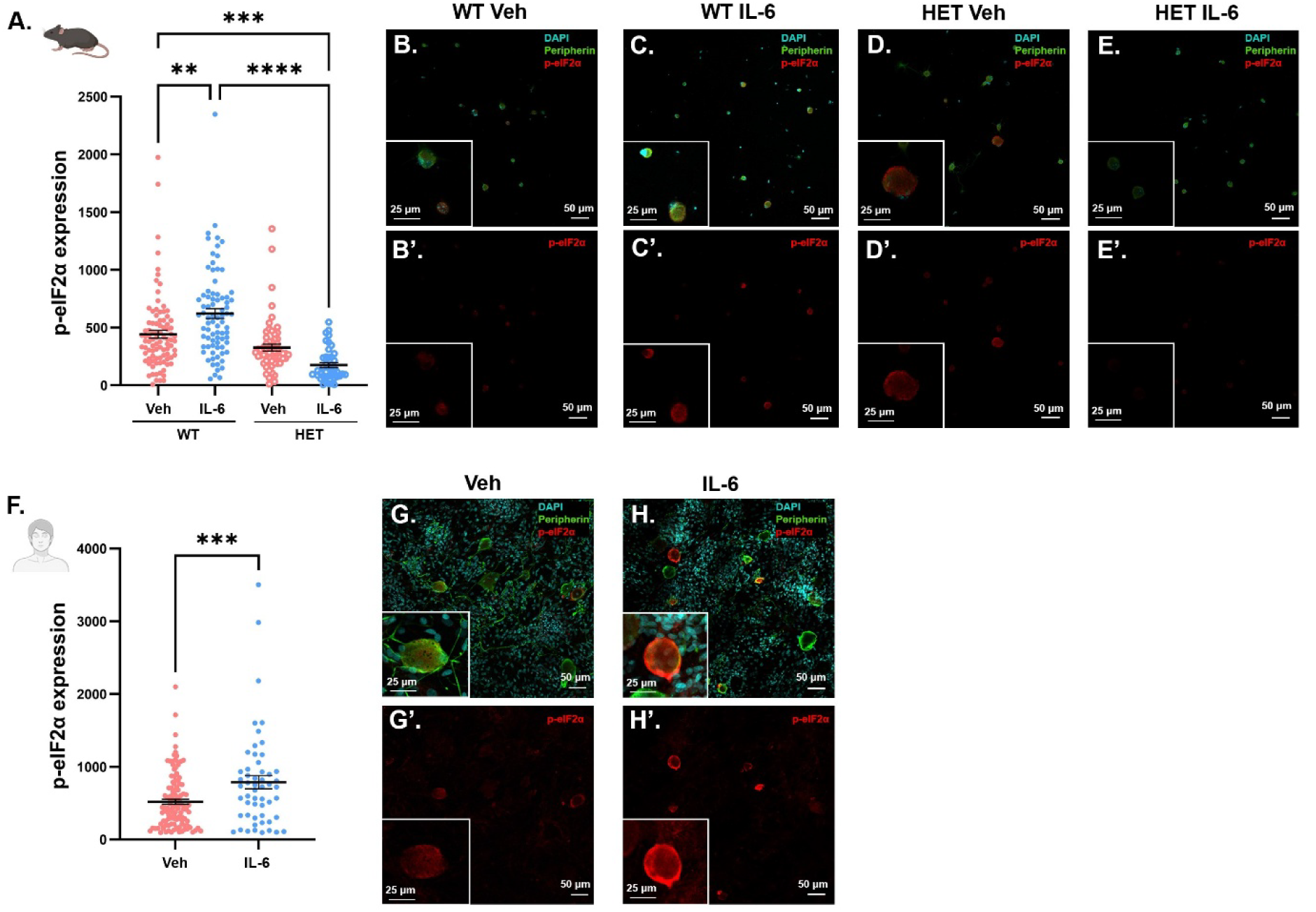
eIF3d Modulates IL-6 Induced p-eIF2α Expression and Integrated Stress Response Activation in Mouse and Human DRG Neurons. (A) p-eIF2α expression is increased between in IL-6 treated WT cultured primary mouse DRG neurons compared to Veh WT neurons (p≤0.01). p-eIF2α expression is unchanged between in IL-6 treated HET cultured primary mouse DRG neurons compared to Veh HET neurons (p>0.05). (B-E’) Representative images of in IL-6 and Veh treated cultured primary mouse neuron from WT and HET mice. DAPI (nuclear marker, blue), Peripherin (neuronal marker, green) and p-eIF2α (red). (F) p-eIF2α expression is increased between in IL-6 treated WT cultured primary human DRG neurons compared to Veh WT neurons (p≤0.001) (G-H’) Representative images of in IL-6 and Veh treated cultured healthy human DRG neurons. DAPI (nuclear marker, blue), Peripherin (neuronal marker, green), and p-eIF2α (red).

### 2.8 Heterozygous eIF3d Knockout Mice Exhibit Reduced Mechanical and Cold Hypersensitivity in the Paw Incision Model

To extend our findings to a model of surgical injury, we employed the paw incision model of postoperative pain, which recapitulates features of local immune activation and cytokine release [30]. To evaluate the contribution of eIF3d to the modulation of this inflammatory nociceptive state, we quantified mechanical and cold sensitivity over a 12 day period following unilateral hind paw incision in WT and HET mice. Compared to WT controls, HET mice demonstrated significantly attenuated mechanical hypersensitivity across all post-injury time points (Figure 6A) (genotype: F(1, 12)=7.748, p=0.0165), with no significant overtime interaction (Figure 6A) (genotype x time: F (2.842, 34.10)=0.6532, p=0.5785). AUC analysis of von Frey time course revealed a significant difference in the disease course of HET and WT mice (Figure 6B) (p=0.0038, unpaired t-test). However, cold allodynia, assessed by acetone response time, was not significantly reduced in HET mice relative to WT (Figure 6C) (genotype: F(1, 13)=1.007, p=0.3338; genotype x time: F (12, 156)=0.4054, p=0.9599). These findings suggest that partial genetic disruption of eIF3d mitigates the development of incision-induced mechanical and cold hypersensitivity, implicating eIF3d as a regulator in the signaling mechanisms that sustain inflammatory pain following paw incision.

**Figure 6.**
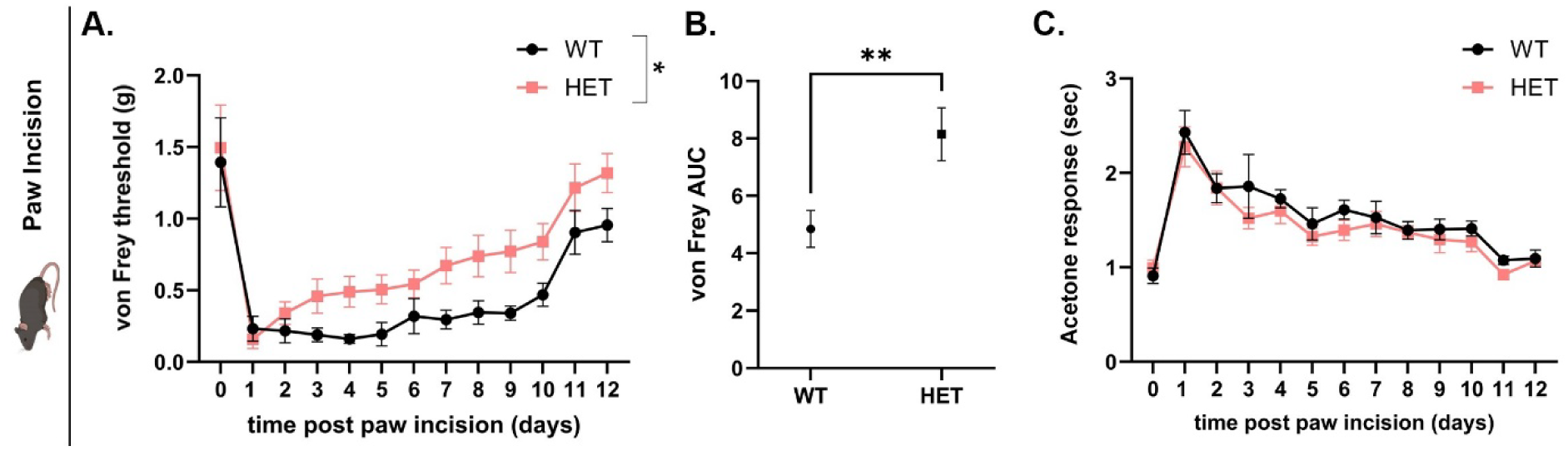
Partial Loss of eIF3d Attenuates Mechanical but not Cold Hypersensitivity Following Paw Incision. WT and HET mice were subjected to hind paw incision, and nociceptive behaviors were assessed. (A) Mechanical hypersensitivity is decreased in HET mice compared to WT mice (p≤0.05). (B) AUC analysis showed significant increase in von Frey threshold indicative of decreased mechanical hypersensitivity (p≤0.01). (C) No difference was observed in the cold hypersensitivity between HET and WT mice as assessed by acetone response time (p>0.05).

### 2.9 Knockdown of eIF3d Does Not Alter Aggregate Pain Scores in the EAE Model of Multiple Sclerosis

Finally, we investigated whether eIF3d modulates the development of pain in the context of neuroinflammation by utilizing the Experimental Autoimmune Encephalomyelitis (EAE) model of multiple sclerosis. Clinical assessments including; disease score (Suppl. Figure 4A) (genotype: F (1, 203)=2.317, p=0.1295; genotype x time: F (28, 203)=0.6163, p=0.9357) and weight change (Suppl. Figure 4B) (genotype: F (1, 175)=0.01691, p=0.8967; genotype x time: F (24, 175)=0.2513, p=0.9999), revealed no significant differences between genotypes or between genotypes over time, indicating that partial loss of eIF3d does not alter susceptibility to or severity of EAE. Maximum disease score attained per mouse each group (Suppl. Figure 4C) (p=0.9176), cumulative disease burden quantified as the aggregate score (Suppl. Figure 4D) (p=0.5186) and the day of disease onset (Suppl. Figure 4E) (p=0.4842) were all unsignificant between HET and WT mice. To evaluate pain related behaviors, we employed nociceptive assays, including mechanical allodynia (von Frey) cold sensitivity (acetone test) and spontaneous pain (mouse grimace scale). Each pain modality was tested at baseline, pre-symptomatic time point and disease onset Across all modalities: von Frey (Suppl. Figure 4F) (baseline: p=0.8833; pre-symptomatic: p=0.9627; onset: p=0.7661), acetone (Suppl. Figure 4G) (baseline: p=0.9164; pre-symptomatic: p=0.9995; onset: p=0.8088), grimace (Suppl. Figure 4H) (baseline: p=0.9763; pre-symptomatic: 0.6187; onset: 0.1014), HET mice exhibited responses comparable to WT controls throughout the disease course. These results suggest that eIF3d is not required for the expression of EAE associated nociceptive behaviors and does not contribute to the development or maintenance of pain in this model.

## 3. Discussion

The overall goal of this study was to define how eIF3d contributes to pain hypersensitivity. We focused on mouse DRGs since DRG neurons are the cellular substrate of ongoing pain behaviors. eIF3d is broadly expressed in human DRGs across neuronal and non-neuronal populations without obvious sex- or size-specific differences. Comparative spatial transcriptomics demonstrate conserved DRG expression across mammals, including rhesus and cynomolgus macaque, guinea pig and mouse, supporting the translational relevance of studying eIF3d in vivo. Accordingly, we investigated eIF3d function using a mouse model, leveraging established genetic tools to probe its contribution to nociceptive processing under physiological and pathophysiological conditions. This conservation supports face validity (eIF3d is present in the same DRG cell classes across species), strengthens translational relevance (mouse results are more likely to reflect human ISR/ATF4/JUN mechanisms) and reduces the risk of a species-specific artifact. We therefore generated and validated a transgenic line, which exhibited substantial reductions in *Eif3d* mRNA transcript and eIF3d protein levels in the DRG, thereby establishing a tractable platform for in vivo mechanistic studies.

At baseline, partial loss of eIF3d did not alter mechanical, thermal, or spontaneous nociceptive thresholds, indicating that normal sensory processing is preserved under uninjured conditions. However, when challenged with pathological stimuli, eIF3d reduction protected HET mice from developing pain hypersensitivity under certain conditions. We and others have previously shown that MGO induces neuropathic pain that is dependent on the stimulation of ISR, such that inhibiting ISR with ISRIB, diroximel fumarate, and FEM-1689, alleviates pain hypersensitivity [5, 31, 32]. In this report, MGO administration attenuated mechanical hypersensitivity in HET mice, particularly at later timepoints, suggesting that eIF3d contributes to the prolongation of MGO-induced ISR, an effect that has previously been reported [11]. Similarly, following IL-6 administration, HETs showed attenuation of mechanical and cold hypersensitivity beginning at 6 hours. A similar protective effect was observed in the paw-incision model, where HETs displayed reduced mechanical hypersensitivity across the postoperative time course. By contrast, eIF3d loss did not alter pain behaviors in the EAE model of T-cell mediated neuroinflammation, suggesting that its contribution is context-specific. The formalin test further emphasized this context dependence: Phase I (peripheral drive) responses were intact in eIF3d knockdown animals, while Phase II (central sensitization) behaviors were modestly increased, pointing to a possible divergence between acute inflammatory signaling and persistent spinal plasticity [33]. The exact mechanisms underlying these context-specific effects require further investigation.

Together, the effects observed in the MGO, IL-6 and incision models point toward a shared mechanism involving ISR. It has been established that incision triggers a robust and sustained upregulation of cytokines, including IL-6, which remain elevated in both tissue and plasma for up to 8 days post injury [30]. In cultured DRG neurons, IL-6 increased phosphorylation of eIF2α, a hallmark of ISR activation, in wild-type cells but this response was blunted in eIF3d-deficient neurons. Behaviorally, this blunting correlated with protection from IL-6 and incision induced hypersensitivity, consistent with the idea that eIF3d facilitates stress-linked translation of pronociceptive effectors. This establishes eIF3d as a context-dependent translational modulator that gates the degree of nociceptive plasticity through ISR-linked pathways in peripheral sensory neurons.

Notably, we observed no difference in hyperalgesic priming between genotypes after IL-6 resolution and subsequent PGE_2_ challenge. Other translation-control pathways, particularly MNK-eIF4E mediated translation, are known to govern and, when inhibited, modulate hyperalgesic priming [34]. This suggests that priming can be maintained via eIF3d-independent mechanisms, with the MNK/eIF4E axis a likely determinant. These findings align with recent work showing that eIF3d is not an essential component of the eIF3 complex in adult tissues, though it appears crucial during embryogenesis, when pluripotent and proliferative cells depend on its translational function [35]. In adult sensory systems, other eIF3 subunits likely compensate, preserving basal translation and nociceptive thresholds. In addition, the set of transcripts it regulates likely varies with cell state and stress modality. For example, Okubo et al. detected no change in JUN translation efficiency [35], whereas in sensory systems JUN/c-Jun is induced downstream of ISR and linked to neuropathic pain [27]. These observations support a model in which eIF3d acts as a selective regulator of stress-adaptive translation, with substrate choice shaped by cellular state and 5′UTR/epitranscriptomic features. From a pain standpoint, heterozygous reduction of eIF3d protects against pain hypersensitivity across multiple inflammatory models.

Our study warrants consideration of several limitations. First, because homozygous deletion is non-viable, we relied on a heterozygous model. As a result, residual eIF3d expression may have masked more profound phenotypic effects. Future investigations employing inducible and/or cell-type specific knockout models of eIF3d could address this limitation by enabling the assessment of complete loss of function in distinct cell types and across timepoints. Furthermore, although eIF3d functions in concert with other subunits to assemble a functional eIF3 initiation complex, we did not evaluate how eIF3d knockdown affects overall eIF3 function or whether other subunits compensate for its reduction. Moreover, while we established that eIF3d knockdown protects against eIF2α phosphorylation in vitro, we have yet to demonstrate this effect in vivo.

Additionally, the specific RNA targets of eIF3d in the dorsal root ganglia remain unidentified. Ribosomal profiling of eIF3d transgenic DRG neurons, for instance, could reveal eIF3d’s influence on translational programming. In summary, eIF3d reduction confers protection from pain hypersensitivity across multiple inflammatory models, likely by limiting stress-responsive translation in sensory neurons. These findings identify eIF3d as a selective, non-essential modulator of nociceptive plasticity, offering a new framework for linking translational control to pain hypersensitivity.

## 4. Materials and Methods

### 4.1 Animals

All animal procedures were conducted in accordance with the Institutional Animal Care and Use Committee (IACUC) guidelines of the University of Texas at Dallas under protocol numbers 14-04 and. This manuscript utilized eIF3d^+/+^ and eIF3d^+/-^mouse lines obtained from Cyagen which were subsequently backcrossed with wild-type C57BL/6 x 129S4/SvJae mice sourced from Charles River. The colony was maintained in-house. The mice used for experiments were between 12 to 24 weeks old. Animals were group-housed under standard laboratory conditions with a 12-hour light/dark cycle and were provided with necessary access to food and water. All efforts were made to minimize animal discomfort and reduce the number of animals used.

### 4.2 Intraplantar Administrations

IL-6 was diluted in PBS to final concentrations of 0.05 ng/μL and 0.02 ng/μL. 10% Formalin (ThermoFisher, 23-245684) was diluted in sterile ddH_2_O to a working concentration of 4%. Methylglyoxal (MGO; Sigma-Aldrich, M0252) was prepared in sterile saline and administered at 20 ng. Intraplantar injections were performed using a Hamilton syringe fitted with a 27-gauge, ½-inch needle. A total volume of 10 μL was delivered into the left hind paw of each mouse.

### 4.3 Paw Incision Model of Post-Operative Inflammatory Pain

The paw incision procedure was performed as previously described [36]. An incision was made through the skin of the plantar surface of the left hind paw, followed by a 2 mm longitudinal cut through the belly of the flexor digitorum brevis muscle. The incision was closed using two 5-0 nylon sutures, placed at each end of the muscle wound. Post-surgical infection prophylaxis was provided via subcutaneous administration of 10% gentamicin (Fisher Scientific, 15-750-060) at a volume of 100 μL.

### 4.4 Behavioral Analysis

#### 4.4.1 von Frey

Animals were acclimated to acrylic rectangular chambers with mesh flooring for 45–60 minutes prior to testing. Mechanical sensitivity was assessed by measuring paw withdrawal thresholds using calibrated von Frey filaments (Ugo Basile, 37450-275) applied to the plantar surface of the left hind paw. Measurements were taken both at baseline and at designated time points following treatment, where applicable. All behavioral assessments were conducted with the experimenter blinded to treatment conditions to minimize bias.

#### 4.4.2 Acetone Test

Mice were habituated to acrylic rectangular chambers with mesh flooring for 45–60 minutes prior to testing. Cold sensitivity was assessed by applying a single drop of acetone to the plantar surface of the left hind paw using a Pasteur pipette. Nocifensive behaviors, including paw shaking and licking, were recorded for 2 minutes following application. Each animal was tested individually and all assessments were conducted under blinded conditions.

#### 4.4.3 Formalin Test

Mice were acclimated to acrylic rectangular chambers with mesh flooring for 30 minutes prior to testing. Animals were briefly removed for intraplantar injection and then immediately returned to the chambers. Nocifensive behaviors as defined by paw shaking, licking and lifting, were recorded continuously for 45 minutes. Each animal was observed individually and all assessments were conducted by an experimenter blinded to treatment conditions.

### 4.5 Mouse DRG extraction and culture

#### 4.5.1 Primary mouse DRG extraction

DRGs were harvested from mice following isoflurane anesthesia and rapid decapitation. The animal was pinned prone onto a Styrofoam platform and the spine was exposed via a dorsal midline incision. Using fine dissection tools under a stereomicroscope, the vertebral column was carefully opened to avoid damaging the spinal cord as to not pull out the DRGs and the spinal cord was displaced to reveal the DRGs. DRGS were collected and cleaned of surrounding connective tissue and transferred to ice-cold HBSS. Approximately 10-15 DRGs were collected per mice.

#### 4.5.2 Primary mouse DRG neuronal cultures

DRGs were harvested from mice and immediately placed in Hanks’ Balanced Salt Solution (HBSS) on ice. Tissue was processed for enzymatic dissociation within 20 minutes of dissection. Ganglia were incubated at 37°C in HBSS containing 2 mg/mL STEMzyme I and 4 μg/mL DNase I in a hot water bath. Following enzymatic digestion, dissociated cells were plated onto poly-D-lysine–coated glass coverslips and maintained in DMEM/F12 + GlutaMAX medium supplemented with 1% Penicillin-Streptomycin and 1% N2 supplement. Cultures were maintained for 2–3 days prior to treatment with 10 ng/mL IL-6/IL-6R complex, diluted in culture media.

### 4.6 DRG Neuronal Cultures

#### 4.6.1 Human DRG neuronal cultures

Human DRGs were obtained from organ donors through the Southwest Transplant Alliance in accordance with published guidelines [37]. Upon collection, culturing process was done according to published protocols [38]. DRGs were immersed in ice-cold artificial cerebrospinal fluid (aCSF) and maintained on ice during dissection. Surrounding connective tissue was carefully removed while submerged in aCSF. Ganglia were minced into ∼2 mm fragments and enzymatically dissociated using a pre-warmed enzyme solution, as described in the referenced protocol. Tissue was subjected to both enzymatic and mechanical dissociation, followed by filtration through a 100 μm cell strainer to obtain a single-cell suspension. Neurons were subsequently enriched using a 10% bovine serum albumin (BSA) density gradient. Cells were plated onto poly-D-lysine–coated glass coverslips in 96-well plates and cultured in DMEM/F12 + GlutaMAX medium supplemented with 1% Penicillin-Streptomycin and 1% N2 supplement. Cultures were maintained for 2–3 days prior to treatment with 10 ng/mL IL-6/IL-6R complex, diluted in the same culture medium.

#### 4.6.2 Immunocytochemistry

Mouse DRG neurons were fixed in 10% formalin for 10 minutes at room temperature, followed by three washes in 1X phosphate-buffered saline (PBS). Cells were then permeabilized and blocked in 10% normal goat serum (NGS) prepared in 1X PBS containing 0.1% Triton X-100. Primary antibodies were diluted in blocking buffer (1X PBS, 0.1% Triton X-100, 2% NGS and 2% fatty acid–free bovine serum albumin) and applied overnight at 4°C. The following primary antibody dilutions were used: DAPI (1:1000), peripherin (1:1000), phospho-eIF2α (1:250) and eIF3d (1:500). Following primary incubation, cells were washed twice with 1X PBS containing 0.1% Tween-20, followed by a final wash in 1X PBS. Secondary antibodies (Alexa Fluor 488 and Alexa Fluor 555) were applied at 1:2000 dilution in blocking buffer and incubated at room temperature for 1 hour. Post-incubation, secondary antibodies were washed using the same protocol as for the primary antibodies. DAPI staining was performed in 1X PBS containing 0.1% Triton X-100 for 5 minutes at room temperature, followed by a single 1X PBS wash. Coverslips were mounted onto glass microscope slides using ProLong Gold Antifade Mountant and imaged using a confocal microscope. Image acquisition and analysis were performed using Olympus CellSens software. Regions of interest (ROIs) were defined around cell somas expressing peripherin and mean gray intensity values were used for quantitative analysis.

#### 4.6.3 Immunohistochemistry

Mouse DRGs were embedded in optimal cutting temperature (OCT) compound and sectioned at a thickness of 12 μm using a Leica CM1950 cryostat. Tissue sections were mounted onto glass slides and fixed in 10% formalin for 10 minutes at room temperature, followed by three washes in 1XPBS using Coplin jars. Sections were blocked in 10% normal goat serum (NGS) diluted in 1XPBS containing 0.1% Triton X-100 for 1 hour at 4°C in a light-protected humidified chamber. Primary antibodies were diluted in blocking buffer (1XPBS, 0.1% Triton X-100, 2% NGS and 2% fatty acid–free bovine serum albumin) and included: DAPI (1:1000), peripherin (1:1000) and eIF3d (1:500). Primary antibody incubation was performed overnight at 4°C in a dark humidified chamber.

Following incubation, slides were washed twice with 1X PBS containing 0.1% Tween-20 and once with 1XPBS for 10 minutes per wash in Coplin jars. Secondary antibodies (Alexa Fluor 488 and Alexa Fluor 555) were applied at 1:2000 dilution in blocking buffer and incubated at room temperature for 1 hour in a dark humidified chamber. Subsequent washing steps mirrored those used for primary antibodies.

DAPI staining was performed by incubating sections in 1XPBS with 0.1% Triton X-100 for 5 minutes at room temperature, followed by a final 1XPBS wash. Sections were mounted with ProLong Gold Antifade Mountant and imaged using a confocal microscope. Image analysis was conducted using Olympus CellSens software. ROIs were drawn around peripherin-positive neurons and mean gray intensity values were used for quantitative assessment.

#### 4.6.4 RT-qPCR

Mouse DRGs were dissected and immediately immersed in Hanks’ Balanced Salt Solution (HBSS). Homogenization was performed in the same collection tubes using 900 μL of QIAzol Lysis Reagent (Qiagen, Cat# 79306). Tissue disruption was carried out with a Minilys homogenizer (Bertin Technologies) at the lowest speed setting using a 30-second burst protocol, conducted in a 4°C cold room to preserve RNA integrity. RNA was extracted using the RNeasy Plus Universal Mini Kit (Qiagen, Cat# 73404), following the manufacturer’s instructions. All steps were performed on ice within a designated RNase-free sterile workspace. RNA concentration and purity were measured using the RNA-specific settings of a NanoDrop 2000 spectrophotometer (ThermoScientific). For cDNA synthesis, reverse transcription was conducted using the iScript Advanced cDNA Synthesis Kit for RT-qPCR (Bio-Rad, Cat# 1725038) per the manufacturer’s protocol. Reactions were carried out on a Bio-Rad T100 thermal cycler. The resulting cDNA was quantified using the single-stranded DNA (ssDNA) mode on the NanoDrop 2000. Quantitative PCR was performed using the SsoAdvanced Universal SYBR Green Supermix (Bio-Rad, Cat# 1725271) on an Applied Biosystems 7500 Real-Time PCR System. Gene-specific RT² qPCR primer assays (Qiagen, Cat# 330001) were used to quantify mRNA expression levels of mouse *eIF3d* (RefSeq: NM_018749). Relative expression was calculated using the comparative CT (ΔΔCT) method and data analysis was conducted using Microsoft Excel and GraphPad Prism.

### 4.7 Data Analysis

cellSens Dimension was utilized for mean grey intensity value and cell diameter analyses. Statistical analysis was performed using GraphPad Prism and Microsoft Excel. Two-way ANOVA with Šídák’s multiple comparisons and unpaired Student’s *t*-tests were used to compare values between WT and HET mice. *P*-values ≤ 0.05 were considered significant, as follows: \**p≤*0.05, \*\**p≤*0.01, \*\*\**p≤*0.001, \*\*\*\**p≤*0.0001, ns = not significant (p>0.05).

## 5. Glossary

ISR: Integrated Stress Response
DRG: Dorsal Root Ganglion
MGO: Methylglyoxal
ROS: Reactive Oxygen Species
AGEs: Advanced Glycation End-products
PGE_2_: prostaglandin E_2_
EAE: Experimental Autoimmune Encephalomyelitis
mTOR: mammalian target of rapamycin
AMPK: AMP-activated protein kinase
MNKs: mitogen-activated protein kinase-interacting kinases

## 6. Acknowledgements

We acknowledge the support of the University of Texas at Dallas Lab Animal Resource Center (LARC) for animal housing and veterinary support. We also acknowledge the Southwest Transplant Alliance (STA) and Theodore Price (University of Texas at Dallas) for their partnership in providing human DRG tissue and thank the donors and their families for their invaluable contributions to this research.

## 7. Author contributions

S.M.M. and S.I.N.: conceptualization, investigation, methodology, formal analysis, data curation, visualization, writing - original draft and writing - review & editing. B.J.W., P.J.P, R.J. A.K., N.E., K.G. and N.A.E.: investigation. M.S.Y.: conceptualization, methodology, investigation, resources, supervision, project administration and funding acquisition

## 11. Figures

**Suppl. Fig. 1.**
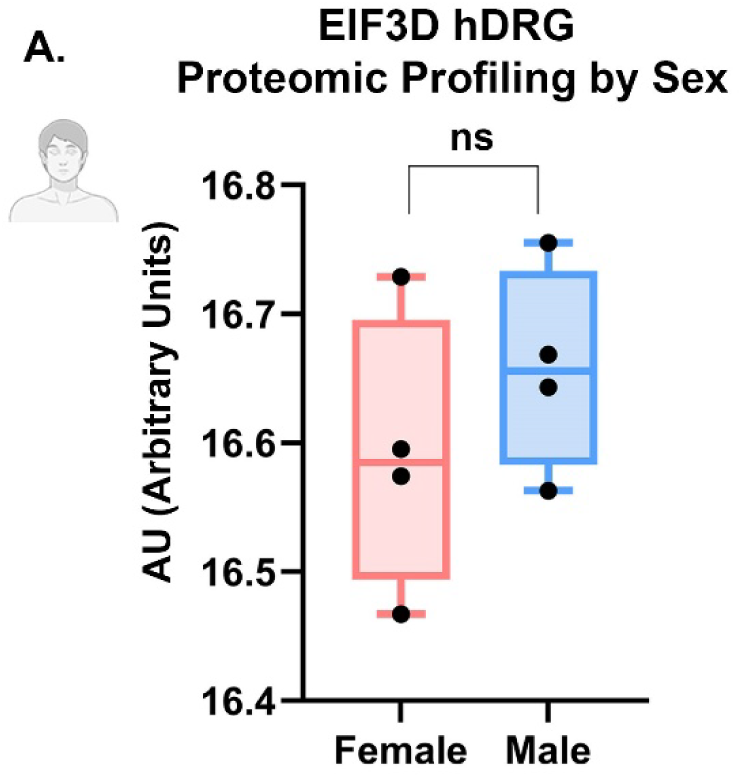
EIF3D Expression in Human Dorsal Root Ganglia Is Consistent Between Males and Females at the Protein Level. (A) Proteomic expression of EIF3D, quantified by arbitrary units (AU), showed no difference between human males and females (p>0.05). *(Data from UTD CAPS Sensoryomics database)*

**Suppl. Fig 2.**
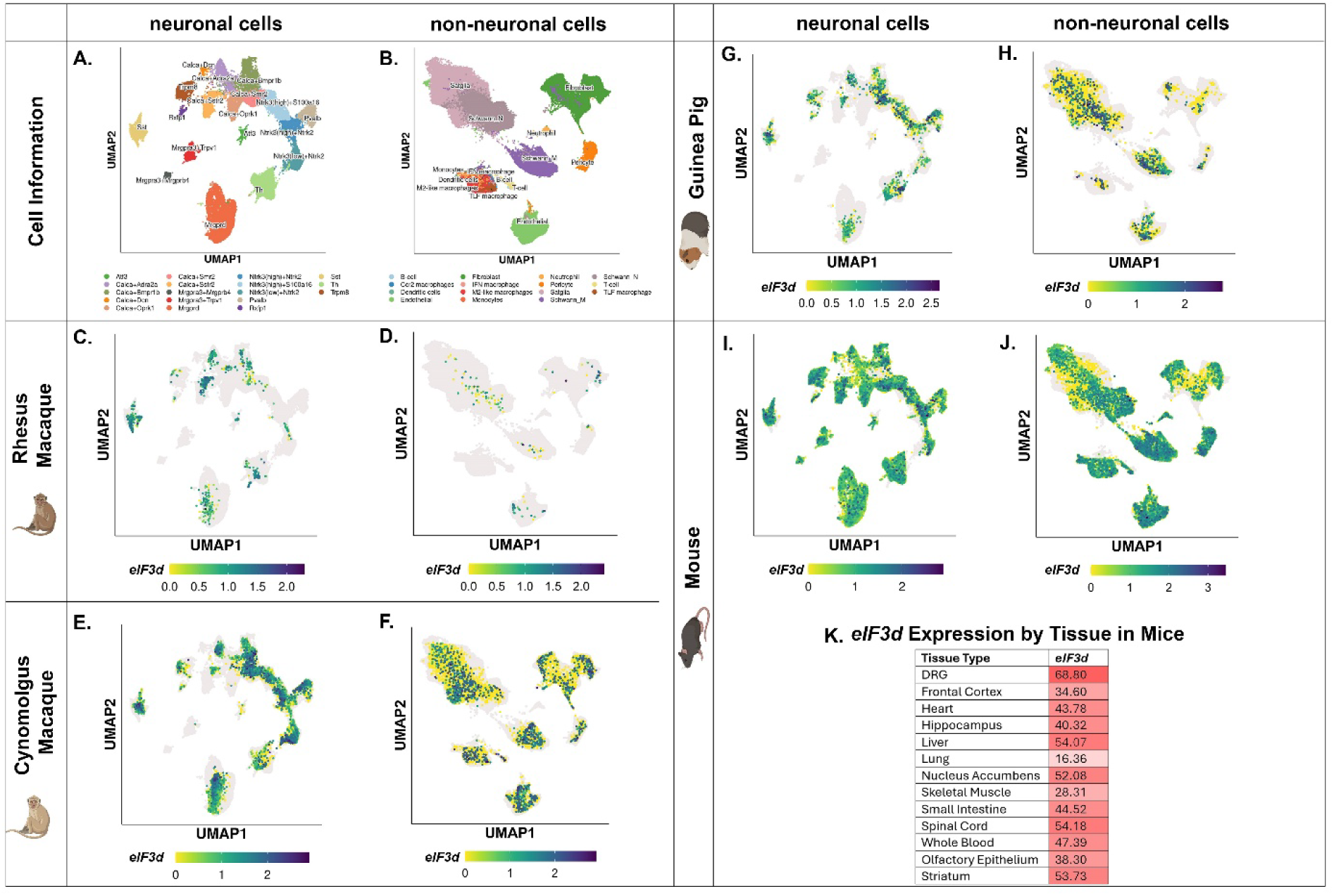
Spatial and Single Nucleus Transcriptomic Dataset Show eIF3d Expression in DRGs Across Mammalian Species, Including but not Limited to Mice. (A) Neuronal cell-type cluster annotations from the spatial and single nucleus transcriptomic dataset. (B) Non-neuronal cell-type cluster annotations from the spatial and single nucleus transcriptomic dataset. (C) *eIF3d* expression in DRG neuronal cells in rhesus macaque. (D) *eIF3d* expression in DRG non-neuronal cells in rhesus macaque. (E) *eIF3d* expression in DRG neuronal cells in cynomolgus macaque. (F) *eIF3d* expression in DRG non-neuronal cells in cynomolgus macaque. (G) *eIF3d* expression in DRG neuronal cells in guinea pig. (H) *eIF3d* expression in DRG non-neuronal cells in guinea pig. (I) *eIF3d* expression in DRG neuronal cells in mouse. (J) *eIF3d* expression in DRG non-neuronal cells in mouse. (K) Transcriptomic profiling of the *eIF3d* gene expression across tissue types in mice, with the highest being in the DRG. *(Data from UTD CAPS Sensoryomics database)*

**Suppl. Figure 3.**
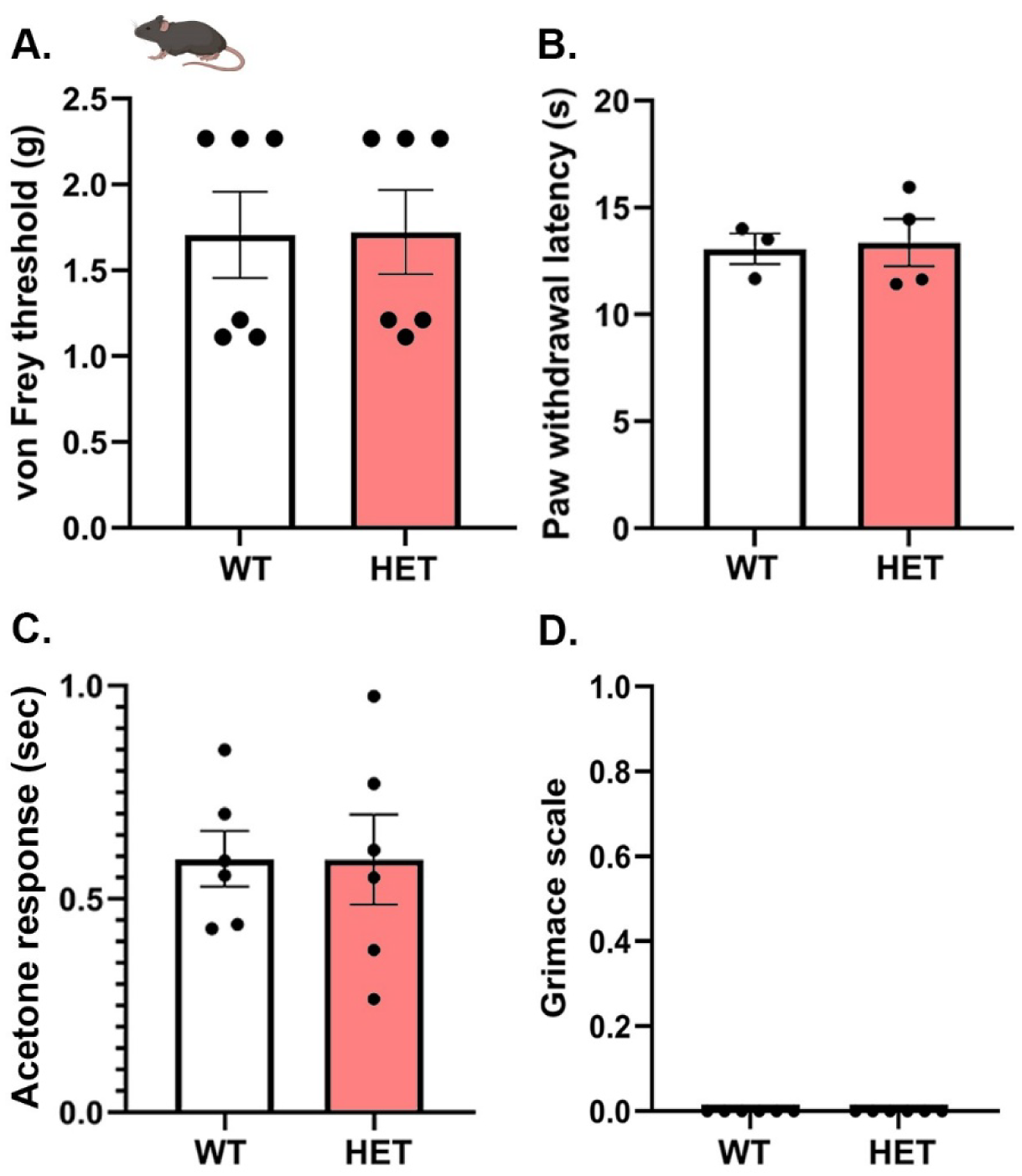
eIF3d Deficient HET Mice Exhibit Comparable Mechanical and Thermal Pain Behaviors to WT Controls. (A) Mechanical sensitivity, tested by von Frey, is not altered between HET and WT mice (p>0.05). (B) Paw withdrawal latency, tested by Hargreaves test, is unchanged between HET and WT mice (p>0.05). (C) Cold hypersensitivity, tested by acetone response time, is unchanged between HET and WT mice (p>0.05). (D) Spontaneous pain behavior, tested by comparison to the grimace scale, showed no pain behavior in either HET or WT mice (p>0.05).

**Suppl. Figure 4.**
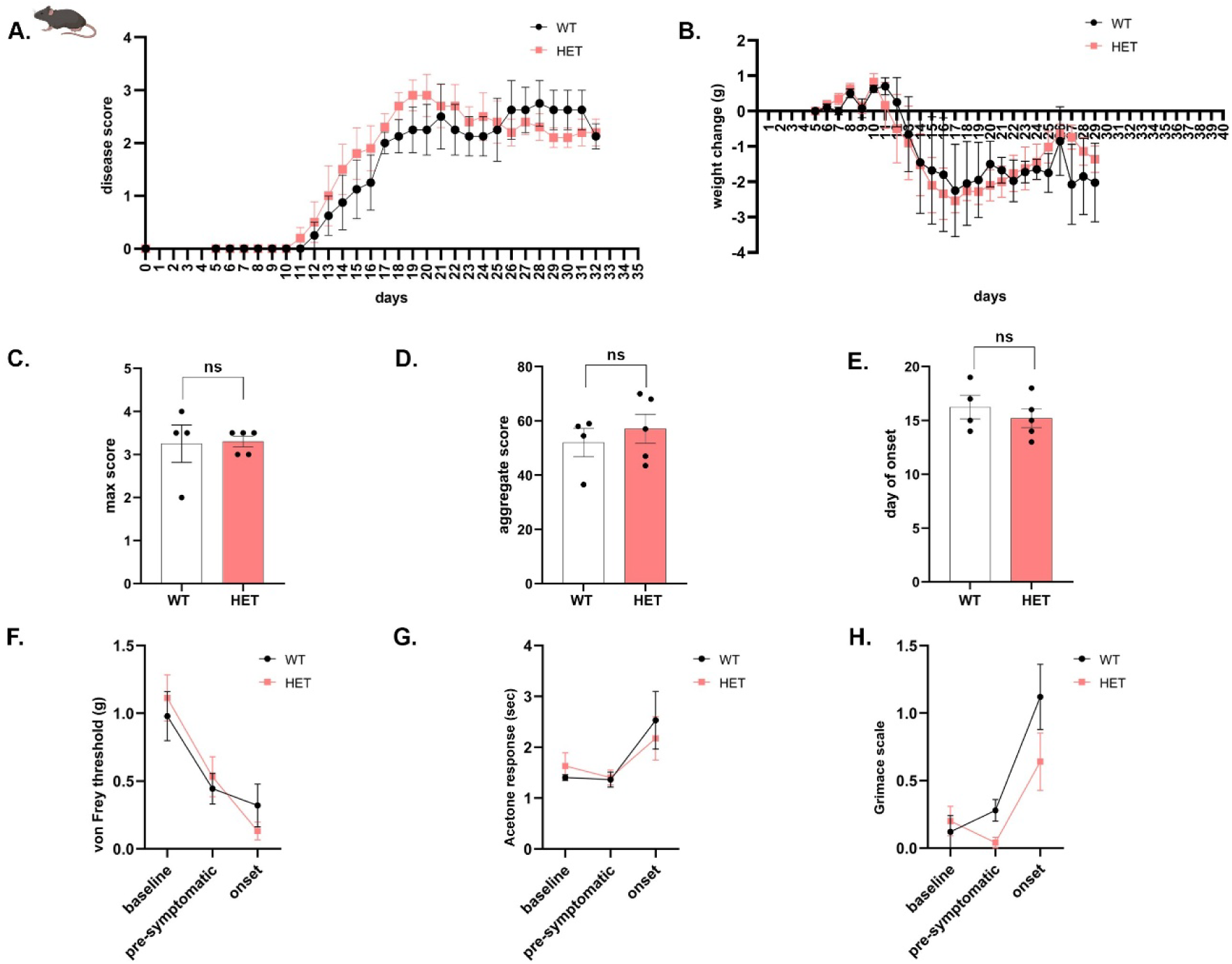
eIF3d Reduction Does Not Affect Pain Behavior or Disease Severity in the EAE Model. (A) Disease progression scores aligned to experimental day 0, comparing HET and WT cohorts (p>0.05). (B) Percent change in body weight over time post-inoculation was unchanged in HET versus WT mice (p>0.05). (C) Maximum disease score attained per mouse in WT and HET groups was unchanged (p>0.05). (D) Cumulative disease burden quantified as the aggregate score between genotypes was unchanged (p>0.05). (E) Onset of disease, expressed as days post-inoculation, was unchanged in WT versus HET mice (p>0.05). (F) Mechanical hypersensitivity, tested by von Frey, at baseline, pre-symptomatic phase, and disease onset was unchanged between genotypes (p>0.05). (G) Cold hypersensitivity, assessed via acetone application, at baseline, pre-symptomatic phase, and disease onset was unchanged between genotypes (p>0.05). (H) Facial grimace scores, an index of non-evoked pain, recorded at baseline, pre-symptomatic timepoint, and disease onset was unchanged between genotypes (p>0.05).

